# Soil microbiome effects on quantitative floral traits of the annual plant *Mimulus guttatus*

**DOI:** 10.64898/2026.10.12.758031

**Authors:** Benjamin A. Sikes, Paige M. Hansen, John Kelly

**Affiliations:** Department of Ecology and Evolutionary Biology, University of Kansas, Lawrence, KS; Kansas Biological Survey and Center for Ecological Research, 2101 Constant Ave. Lawrence, KS; Department of Soil and Crop Science, Colorado State University, Fort Collins, CO, USA

**Keywords:** microbiome, Mimulus, flowering traits, plant-soil feedbacks, rhizosphere

## Abstract

Plant flowering is a critical determinant of fitness and supports pollination services to ecosystems. Flowers are energetically expensive and require nutrients that plants might otherwise use for growth. Soil microorganisms can alter plant development in a variety of ways, but the quantitative importance of microbiomes in determining floral variation has yet to be established. In particular, it is unclear the extent to which genotypic differences in flowering phenotypes are mediated through the microbiome. We used a two-generation plant-soil feedback design to test whether flowering traits in a short-lived annual plant *Mimulus guttatus* drive differentiation in rhizosphere microbiomes. Second, we tested whether these microbiomes subsequently affected flowering traits of progeny plants. We found that different plant genotypes did not generate differences in rhizosphere microbiome structure or diversity after eight weeks of growth. When these microbiomes were inoculated onto new plants, flowering traits were mainly driven by the progeny plant’s genotype and not the genotype of the parental plant that trained the inoculated microbiome. Longer microbial training, as would necessarily happen in perennial plants, may produce stronger effects, but our results suggest microbiome effects contribute mainly to the “environmental noise” in flower traits in the short-lived annual *Mimulus guttatus*.

## INTRODUCTION

Flowering phenotype is a major determinant of plant fitness (Schemske and Bradshaw Jr, 1999; Brown and Kelly, 2018; Gaudinier and Blackman, 2020) and the ecosystem services that plants provide (Campbell et al., 2012; Lowe et al., 2021). Flowers are energetically expensive and pollinators may only be present at specific times with visitation and successful outcrossing depending on floral traits such as display size and flower number (Goodwillie et al., 2010). Apart from genetic factors, multiple environmental factors can alter flower traits (Reeves and Coupland, 2000; Cho et al., 2017) and anthropogenic climate change is shifting both the timing of flowering and flower morphology (Bishop et al., 2023; Acoca Pidolle et al., 2024). Understanding the genotypic and environmental forces that shape flowering phenotypes is critical for both basic plant biology and to select for flowering traits that best optimize these services, particularly with increasing environmental change (Fitter and Fitter, 2002). Successful pollination is critical both to maintain wild plant populations and for pollination-dependent crops.

The effect of microbiomes on flowering traits is an understudied aspect of plant biology. Many different plant traits shape the structure and function of their microbiomes, particularly rhizosphere microbiomes (Wagner et al., 2016; Wagner, 2021). Artificial selection on plant flower timing in *Arabidopsis thaliana* has been shown to alter rhizosphere microbiomes and, in turn, inoculation of these microbes shifts the flowering of naïve hosts (Panke-Buisse et al., 2015; Lu et al., 2018). This suggests microbiome composition, and its effects on plant performance, can be specific to the genotypes segregating in a population. When this happens, the transmission of microbiomes from parents to offspring becomes an important determinant of trait variation and heritability (Henry et al., 2021; Week et al., 2025). Experiments testing these feedbacks, where parental plants “train” a microbiome that is subsequently transferred to and may effect offspring plants, are commonly used to measure microbial effects on the species composition of plant communities (Bever et al., 2012; Van der Putten et al., 2013). There has been less exploration of feedback effects among genotypes within a single species on traits. The critical question is whether differentiation of rhizosphere microbes is an important source of flower trait variation. If so, they might practically be utilized to optimize plant fitness and ecosystem services.

Our experiment employed a two-step plant-soil feedback design (Figure 1) using inbred lines of *Mimulus guttatus* (yellow monkeyflower) derived from a natural population in Oregon. The genetics of floral traits have been thoroughly studied in *Mimulus* but little is known about their microbiomes (Bradshaw Jr et al., 1995; Wu et al., 2008; Monnahan and Kelly, 2017; McIntosh et al., 2024). We examined genotypes from the Iron Mountain population of *M. guttatus* where flower size related traits exhibit high variability with 30-40% of the variation attributable to additive genetic effects (Kelly and Arathi, 2003). For this experiment, we selected two homozygous lines with distinct flowering phenotypes (Troth et al., 2018). The IM62 genotype is slow to flower but produces large flowers. The IM767 genotype progresses to flower rapidly but then produces smaller flowers. In the “training” generation of our experiment, these genotypes were grown in a diverse, but naïve-to-*Mimulus* soil microbiome until flowering. We used amplicon sequencing to explore microbiome differentiation derived from the training step genotype. In the second generation, each genotype was grown in a common sterile background soil inoculated with microbes derived from one of the two training genotypes or with a sterilized inoculum. If the training generation generates a microbiome specific to each genotype (either compositionally or functionally), we predict altered flowering traits *within* each genotype contingent on which parental genotype conditioned their inoculum. In contrast, if microbiomes are similar for parental genotypes, we may see a difference between live and sterilized inoculums (either positive or negative), but no interaction between parental-soil-training genotype and offspring genotype in determining offspring phenotypes. This work advances our understanding of the speed with which short-lived plants can alter microbiomes and whether these can influence fundamental traits for plant fitness, pollinators, and the food supply that depends on flowering plants.

**Figure 1:**
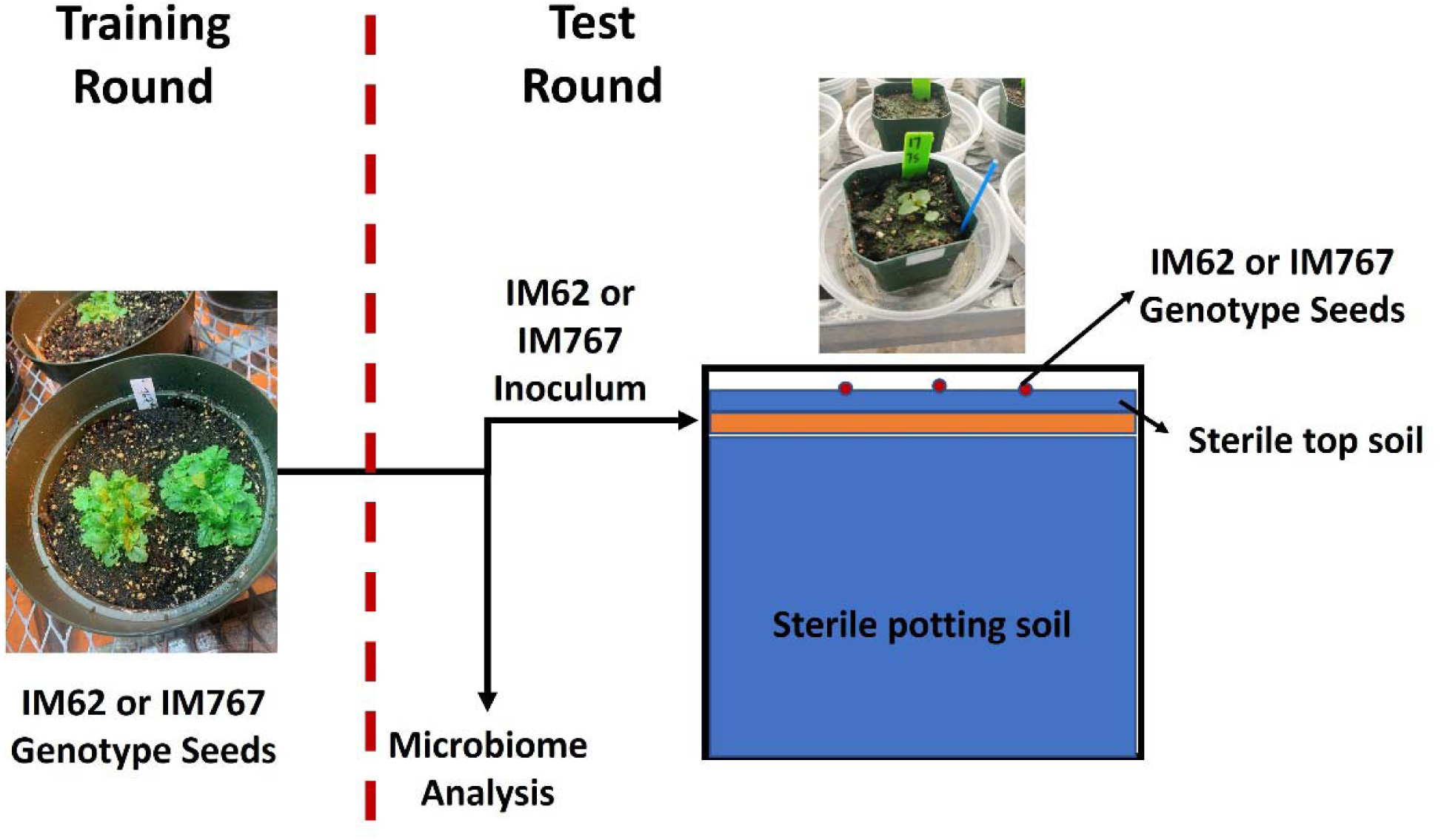
Experimental design to test feedbacks between genotype-based flowering traits and rhizosphere microbiome.

## MATERIALS AND METHODS

*Mimulus guttatus* (syn *Erythranthe guttata*; 2n = 28; Phrymaceae) is a model plant for genetics and mating systems evolution (Wu et al., 2008). The genotypes used in this experiment were sampled from Iron Mountain in Oregon (IM; 44.402217N, 122.153317W; (Willis, 1999; Kelly, 2003). This population reproduces primary by outcrossing at an average rate of about 90% (Willis, 1993; Monnahan et al., 2021). Starting with seed from thousands of unrelated field plants, inbred lines were created by successive rounds of self-fertilization, a.k.a. single seed descent (Willis, 1999; Kelly, 2003). We selected IM62 and IM767 from this large collection of inbred lines, all of which have been characterized for 13 flower size and developmental timing traits known to influence fitness in the field (Troth et al., 2018). These data confirm that IM767 (small flowers, fast development) and IM62 (large flowers, slow development) are divergent with respect to the tradeoff between rapid development and large flower size. Several key traits underpin this tradeoff and are easily measurable, including day to first flower, pistil length, and corolla width.

### Experimental design

Standard potting soil was sterilized to remove existing microbes via two rounds of autoclaving at 121°C for 1 hour each, with a 24-hour rest between. We then mixed this soil in a 50:50 ratio with field soil from local prairie and agricultural sites to present a diverse starting microbiome on which the different genotypes could select. This starting inoculum includes microorganisms evolved with perennial and annual wild plants (grass, forbs, and legumes) as well as those under considerable selection for crop production, mainly a corn/soybean rotation. However, none of the sites include *M. guttatus*. Into these soils we planted seeds from one of the two *M. guttatus* inbred lines (IM62 or IM767). In the “Training Round” (Fig 1), one-gallon pots (25cm in diameter) pots were filled with the potting/field soil mix for microbiome selection and 10 seeds of each plant were added to each pot. To maximize potential microbiome components that could be selected on, field soils were sourced from two post-agricultural sites and a native prairie site at the KU Field Station. Twenty individual training pots were created for each genotype (40 total). Plants were grown in a greenhouse in October and November under a 14hr light/10hr dark cycle, with supplemental lighting provided by halogen lamps. Soils were moist at the start and pots were bottom watered using sterile, ultra-pure water every 3 days to ensure relatively wet soils. This watering is ideal for *M. guttatus* growth to simulate their brief life cycle following snow melt (personal observation). Plants flowered 4-6 weeks after seeding depending on genotype, and all watering was stopped after 6 weeks allowing plants to begin senescence and soils to dry out. On November 19^th^, we harvested the training round, noting the # of individual plants and their total aboveground dry biomass, following plant biomass drying in an oven for 3 days at 110°C. We harvested the upper 5cm of soil where roots were most concentrated (i.e., rhizosphere). Soils from each training round pot were kept separate in sterile Ziploc bags, homogenized thoroughly, and a 5% subsample (∼20g) of each was taken for DNA analysis and placed in a -80°C freezer. Training round soils were kept at 4°C and used as inoculum for the “Test Round” (see below) within 2 weeks.

### Microbiome Analyses

We used 16S amplicon sequencing to map microbiome changes between divergent flowering genotypes in the training round. We focused on bacteria because their populations grow more rapidly than fungi and other soil biota (Koch et al., 2018; Martinović et al., 2021). As such, their communities are more likely to respond to plant trait differences in a single generation, particularly for a short-lived plant. In addition, bacteria generally grow more rapidly and inhibit fungi in wet soils (Hicks et al., 2019) like those in which *Mimulus* were grown. DNA was extracted from 0.25 g of subsampled soil using MoBio PowerSoil Kits (MoBio, Carlsbad, USA). Extracted DNA was quantified using Qubit 2.0 (LifeTechnologies, Carlsbad, USA) then the V4 hyper-variable region of 16S rDNA was amplified from 5 ng of template DNA using the standard Earth Microbiome Primers 515F and 806R (Caporaso et al., 2018) and Q5 polymerase (New England Biosystems,Ipswich, USA). The PCR protocol consisted of an initial denaturing at 98 °C for 30 s; then 25 cycles of denaturing at 98 °C for 10 s, annealing at 55 °C for 30 s, and extension at 72 °C for 30 s; followed by a final extension at 72 °C for 5 min. PCR products were cleaned using Agencourt AMPure XP magnetic beads (Beckman Coulter, Indianapolis,USA). A second PCR was then used to ligate unique Nexterra indices (Illumina, San Diego, USA) to each sample. Cycling conditions for the indexing PCR were similar to the first PCR, except for only running for 8 cycles, and then bead purified as before. Individual libraries were pooled at equimolar concentrations into a single, 4 mM library; concentration and amplicon size was verified using a TapeStation 2200 (Agilent, Santa Clara, USA). Amplicon sequences were then generated using a 301 bp, paired-end run on an Illumina Mi-seq (Illumina, San Diego, USA) at the University of Kansas Genome Sequencing Core. Sequencing data included a negative water-only library control (from 1st PCR step above).

### Bioinformatics

We used a standard Qiime2 bioinformatics pipeline (Bolyen et al., 2019) to produce a table of amplicon sequence variants (ASVs) that we could compare depending on the *Mimulus* genotype training. Demultiplexed paired reads were denoised, merged, and chimera-checked using the DADA2 implementation in Qiime2 v9 (2024.10). The ASV table was imported into R, then we used phyloseq (McMurdie and Holmes, 2013) and related packages to complete bioinformatics and statistical analysis. We assigned taxonomy for individual ASVs using alignment to the SILVA database (Glöckner et al., 2017); v. silva_nr99_v138.1). We carried out additional quality filtering to remove any Eukaryotic taxa and filter failed samples (<1000 reads based on water negative control). We also focused on dominant taxa by removing any ASVs that occur less than 3 times and 3 samples as sequencing artifacts. For differential abundance of individual taxa based on Genotype, we used the DESeq2 package (Love et al., 2014).

### Data analysis

For alpha diversity, we used general linear models (glm) to explore the effect of Genotype or Plant Biomass on the three Hill components (raw, Shannon, Inverse Simpson) and Faith’s Phylogenetic Diversity. We included read depth in each model to account for variations in sequencing depth. For beta-diversity, we used compositional data analysis to determine community differences (Quinn et al., 2019), transforming the data using a center log-ratio and using Aitchison’s distance to calculate sample dissimilarity. We used PERMANOVA in the adonis2 implementation in the vegan package (Oksanen et al., 2013) to contrast community differences based on training genotype and biomass. Finally, DESeq2 was used to identify differentially abundant taxa among training genotype. Full scripts for bioinformatics and statistical analyses are available upon request.

### Test round experimental design, data collection, and analysis

A second generation, the “test-round” (Figure 1), was set up in sterilized 6.25 cm^2^ pots. We used a sterile base of 50:50 potting soil/sand mix, sterilized twice via autoclaving as before the training round. Training round soils were kept distinct by pot so we could carry through and estimate among pot variation. Soils from each training round pot were homogenized and 50ml used as a thin inoculum layer (∼5% of pot volume) sandwiched between two sterile soil layers (see Figure 1). We had four inoculum treatments in the test round: IM62 inoculum, IM767 inoculum, or sterilized version of each of these to account for any abiotic effects. Live or sterile inocula were blocked by training pot. Into these, three seeds of one of the two *Mimulus* genotypes were sown. After germination, plants were thinned to a single individual that was most dominant.

In total, each test genotype in each inocula treatment was replicated 20 times, blocked by maternal pot for a total of 160 pots. *Mimulus* did not germinate in 19 pots, and plants in 46 additional pots died after germination before any flowering data could be collected. For the remaining plants we collected data on date of first flower, pistil length and corolla width for the first flower, and aboveground biomass at harvest at the end of the 2^nd^ growth period. We used mixed effect models with Type III ANOVAs to test for differences in the test round for the four plant response variables based on test plant genotype, the genotype that trained soils in the first round (hereafter maternal genotype), and whether the soil was alive or dead. Training generation pot was nested within maternal genotype to further account for variation from this round. We assessed these differences using R (R Core Development Team, 2022), including the lme4 package (Bates et al., 2015) for mixed effect models, ggplot2 (Wickham, 2016) for graphics, and the effectsize package (Ben-Shachar et al., 2020).

To test whether specific microbiome species transferred in live inocula affected plant traits, we used a regularized regression approach. Specifically, we applied elastic net regression (Friedman et al., 2010; Tay et al., 2023) to predict the four trait values measured in round 2 (days to flower, flower width, pistil length, and biomass) among the plants receiving a live inoculum. Individual genotype (IM767 or IM62) was included as a binary predictor, as were the abundances of each taxon identified from 16S amplicon sequencing after quality filtering. Given that the number of predictors is much greater than the number of observations, elastic net regression uses a combination of Lasso and Ridge penalizations to prevent overfitting. The two parameters that determine the relative importance of Lasso and Ridge, α and λ, are calculated using a stochastic cross-validation process. We replicated this stochastic process to calculate an average α and λ for each trait. The final regression model applied to each trait (based on these α and λ values) is deterministic. These calculations were conducted using the “glmnet” library of R (Friedman et al., 2023).

## RESULTS

### Training Round and Microbiome analysis

Plant genotype in the training round followed expected genotypic differences: IM62 plants flowered later, producing flowers with wider corollas and longer pistils than IM767 plants. The biomass of IM62 plants was greater than IM767 when measured after 60 days. Microbial diversity and community composition were not affected by training round genotype. Genotype did not alter alpha diversity across Hill numbers representing ASV richness (LR Chisq_1,35_ = 0.000, P = 0.989), Shannon (LR Chisq_1,35_ = 0.140, P = 0.709), and Inverse Simpson’s diversity (LR Chisq_1,35_ = 0.504, P = 0.478), which differ in their weighting of rare vs dominant taxa. Increasing host biomass tended to decrease phylogenetic diversity (Faith’s PD; LR Chisq_1,35_ = 0.977, P = 0.323; Supplemental Figure 1), but neither biomass nor training genotype were significant predictors. Variation in community composition was not based on the genotype of the training plants (F_1,36_ = 0.9934, P = 0.545, Figure 2), the total biomass of plants in the training round (F_1,36_ = 0.9858, P = 0.624), or their interaction (F_1,36_ = 0.9521, P = 0.843). Four taxa were differentially abundant (adj.P <0.05) based on training Genotype. In microbiomes trained by IM767, an amplicon sequence variant (ASV) in the genus Luedemannella (Actinobacteriota; Micromonosporaceae) had a 23.5 log2fold increase and an ASV in the family Entotheonellaceae (Entotheonellaeota) had a 7.3 log2fold increase compared to IM62 microbiomes. In IM62 trained microbiomes, an ASV in the order Solirubrobacterales (67-14 family) and an ASV in the phylum Chloroflexi (TK-10 class) had log2fold increases of 22.5 and 22.3 respectively, compared to IM767 trained microbiomes.

**Figure 2:**
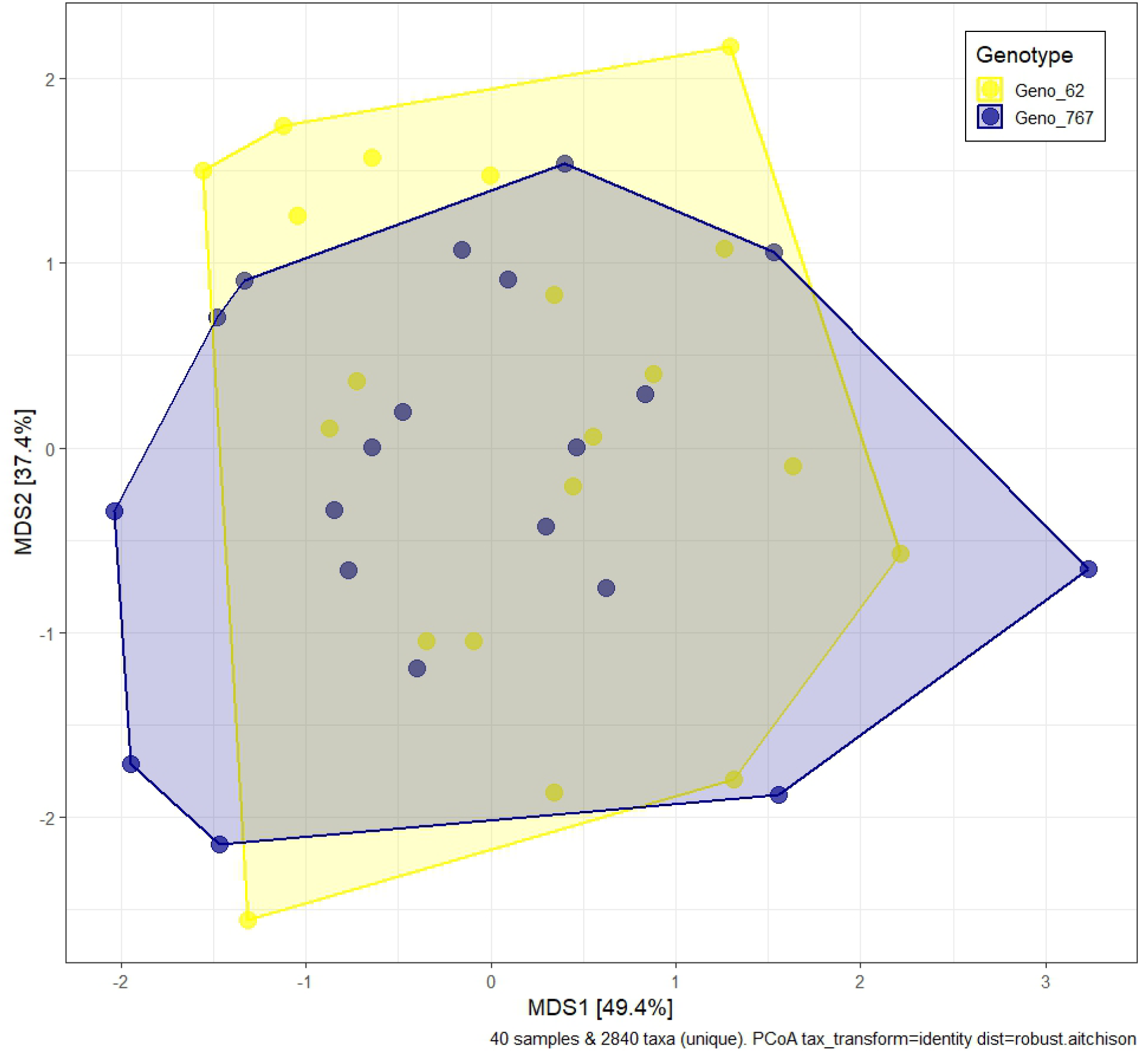
Microbiome composition between training round genotypes. First and second axes of principle coordinate analysis are based on center log-ratio transformed data. Proportional variation explained is given with each axis and training round genotype follows colors in legend — blue for IM767 and yellow for IM62.

### Test Round

Plant genotype was the dominant effect for all plant traits measured in the test round, explaining between one-third and one-half of variation in time to flower, flower size, pistil length, and biomass. Plant genotype effects independent of microbial effects significantly predicted time to flower (F_1,65.1_ = 55.19, P <0.001), corolla width (F_1,75_ = 66.52, P <0.001), and biomass (F_1,75.5_ = 105.74, P <0.001). Plant genotype strongly predicted pistil length (F_1,76_ = 47.45, P <0.001) but also had a higher order interaction with live/sterile and the maternal genotype that trained the inoculum (3-way: F_1,76_ = 4.52, P <0.05). Plants from the IM62 genotype, on average, produced 4.5x more biomass, took 10 days longer to flower, and produced flowers with 4.45mm (22.6%) larger corollas and 1.33mm longer pistils (9%) than IM767 plants (P <0.05 for all).

Inoculum had weaker but additional effects on plant flowering traits. For time to flower, the interaction of maternal genotype and live/sterile inoculum explained 7% of proportional variation (F_1,67.6_ = 5.31, P <0.05). Compared to IM62-trained microbes, plants inoculated with microbes trained by the IM767 maternal genotype (fast flowering) had slower flowering for both test plant genotypes, on average by about 5.5 days. Adding sterilized inoculum (regardless of training) made plants flower similar to those inoculated with IM62-trained soils (Figure 3). Biomass was strongly correlated with days to first flower (0.771; Figure S1), but with weaker inoculum effects (Table S1): the interaction between maternal genotype and live/sterile soil was marginally significant (F_1,74.9_ = 3.10, P =0.082; partial η² = 0.04). Floral morphology showed more variation from inocula. Plants grown in live vs. sterile soil differed in flower width (F_1,75_ = 6.29, P <0.05; partial η² = 0.08, Figure S2a), while pistil length showed a more complex relationship to inoculum, even though it was 75% correlated with corolla width (Table S1, Figure S3). For pistil, interactions among genotype, maternal microbial training, and live/sterile (see above) were largely genotype-specific response to soil sterilization (Rd2.Genotype:Live/Sterile: F_1,76_ = 10.33, P <0.005, partial η² = 0.12). IM767 plants made longer pistils in sterile than live soil, but without a similar effect for IM62 plants. The 3-way interaction (noted above, partial η² = 0.08) showed live IM767-trained soil increased IM62 test plant pistil lengths over sterile, but decreased pistil length of IM767 test plants.

**Figure 3:**
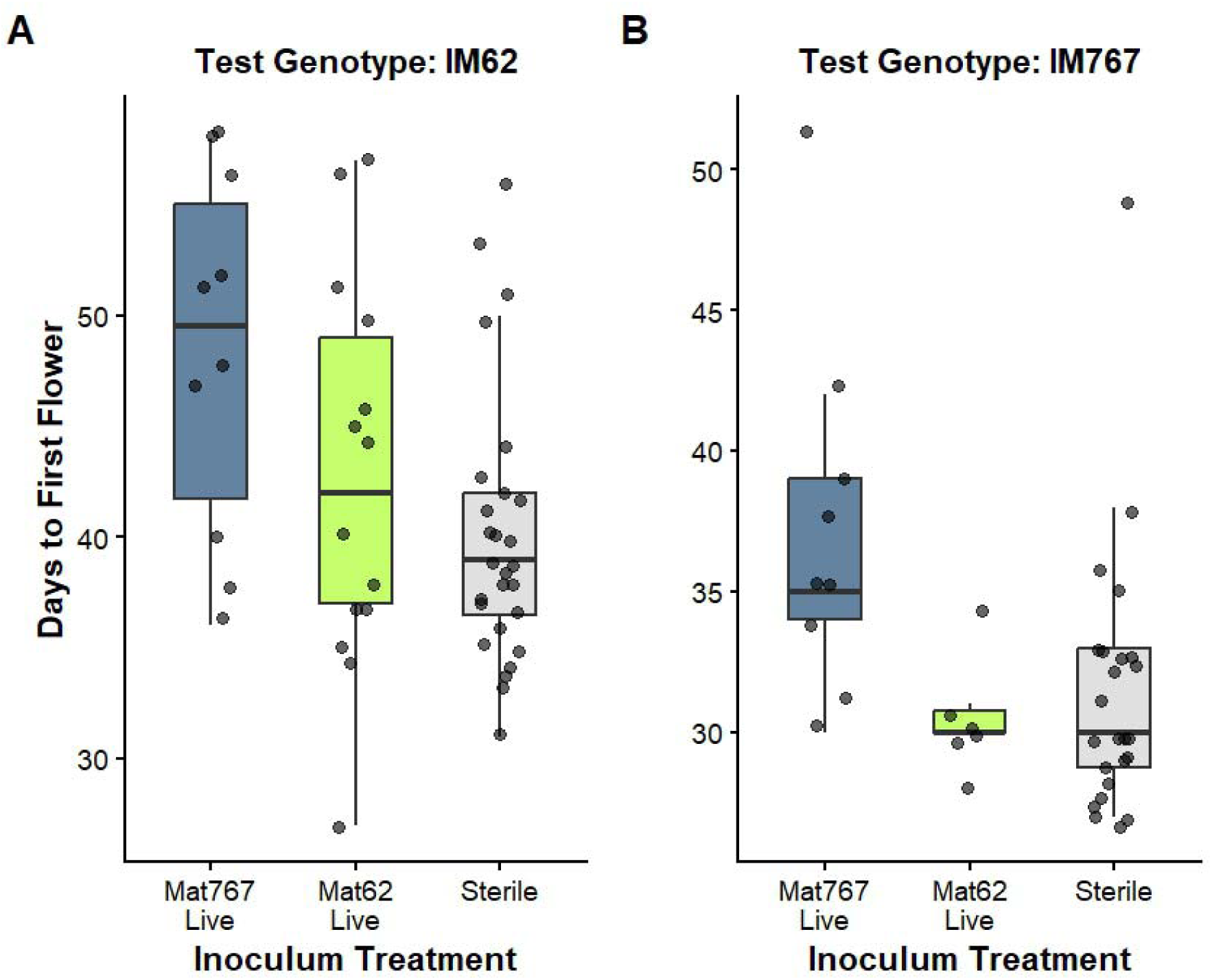
Days to First Flower response to test plant genotype, panel A = IM62, panel B= IM767, training plant genotype for microbiome, or sterile inoculum. X-axis labels and colors represent the two maternal genotypes: Mat767 (dark blue) or Mat62 (chartreuse), or sterile (gray) inocula. Box and whisker plots indicate median (line), 75% of data (boxes), and 95% (whiskers).

Regularized regression showed that several microbial taxa from the training round significantly predicted traits in Round 2, with days to flower and flower width both being well predicted (Table 1). The abundance of an aerobic, gram-negative Acetobacteraceae was associated with longer flowering time, a nearly 60% increase from that associated with Rd2 genotype. Accounting for taxa abundance, genotype has the fourth strongest effect on flower width, with the three more correlated taxa all associated with much narrower flowers. Most taxa whose abundance predicted large shifts in one trait did not have large effects on other traits as well. Supplementary Table S2 shows the top 5 elements by coefficient for each trait in Round 2, Figure S4 shows a heatmap of these ASV elements across the 4 traits, and Table S3 shows the seventeen taxa whose abundance predicted two or more traits.

**Table 1.** Model summaries from regularized regression of plant traits onto microbiome abundances contributed by parental pots.

| Trait | Number of significant predictors | R<sup>2</sup> |
| --- | --- | --- |
| Days to flower | 30 | 0.92 |
| Flower width | 46 | 0.93 |
| Pistil length | 23 | 0.71 |
| Biomass | 32 | 0.85 |

## Discussion

Genotype was the dominant influence on flowering traits for a short-lived annual plant. Despite manipulation of host microbiome and multiple rounds of growth in the greenhouse, flowering trait differences were explained mainly by genotype (nearly 50% of the overall variance). The two genotypes examined here, IM62 and IM767, were derived from one natural population (Willis, 1999) but selected from a large panel of such lines based on previously established differences in phenology and flower morphology (Kelly and Arathi, 2003). Microbiome variation may have contributed significantly to the “environmental variance” in traits, a contention supported by the significant prediction of plant phenotypes from microbiome community abundances within inocula (Table 1). However, microbiome responses were not strongly genotype specific (Fig 2, Table S1).

Microbiome influences on traits like flowering likely depend on plant life history or continued selection. A single round of growth for a short-lived annual plant was not enough to significantly shift microbiomes, either in diversity, community composition, or their functional effect on flowering in the successive generation. Longer, consistent selection by the same genotype, either through successive generations or perennial host plants, may produce microbiome differences like those seen in previous experiments (Panke-Busse et al. 2015) and be more likely in the field. Greater microbiome differences could drive greater effects on flowering and beyond. However, shifts in the rhizosphere microbiome may be more likely driven by root traits, like architecture or exudation (Wen et al., 2025), compared to flowering traits. Eliminating the rhizosphere microbiome (i.e., sterilization) had effects on all three flowering traits (either alone or through interactions). These generally resulted in faster flowering, and larger flowers with longer pistils, traits that summarize a phenological tradeoff for plants in the field. These shifts may be particularly important for plants like *Mimulus* that must time their growth and reproduction to take advantage of the short warm, wet period after snow melt (Hall and Willis, 2006).

Despite not showing strong genotype effects on microbiomes, plant traits did still respond to microbiomes. Although not a component of our hypotheses, microbiome effects were common when contrasting live vs. sterilized inoculum and these effects sometimes differed with the maternal genotype which trained the inoculum. Inoculation with live soil delayed progression to flowering relative to sterilized soil (p<0.006, Table S1). Here, there was a subtle genotype dependency because the magnitude of the delay differed between soils trained by the two genotypes (p=0.02, Table S1, Fig 3). These results parallel findings from another recent study of *M. guttatus*: McIntosh et al (2024) measured plant growth and development with both live and sterilized inoculums. While their study considered genotypes sampled from different habitats, these authors also found that plants flowered more rapidly when the inoculum was sterilized. All else equal, more rapid flowering should be favorable, and thus these results suggest that genotype specificity of natural enemies within the microbiome might be an important selective agent in this species.

Microbial taxa identified here show two important trends related to flowering traits that might differ with repeated selection. In our study, the abundance of specific microbial taxa was clearly associated with maternal genotype, even without broad shifts in microbiome diversity and composition. However, the microbiome taxa with the strongest associations with trait variation in Round 2 (Table S2) were not those that showed the greatest differentiation between IM62 and IM767 parents (the training environment of Round 1). Regularized regression showed that individual taxa transmitted from parent plants have both positive and negative effects on plant traits. Because these taxa are not determined by the parental generation genotype, they are a component of the environmental variance when considered from a quantitative genetic perspective. Transmissible taxa that do not shift with genotype could be ideal for applications in agriculture (Arif et al., 2020) and ecosystem restoration (Robinson et al., 2023). Interestingly, repeated rounds of selection tend to produce host-independent effects following several rounds (Morella et al., 2020), in contrast to a single round tested here. Repeated genotypic selection, which may be common in the field, may have distinct outcomes on the degree of microbial shifts, their flowering effects, and their relative importance compared to host genotype.

The plant-soil feedback (PSF) design should be more widely used to test microbial contributions to traits critical for plant fitness, productivity, and function. The ecological theory and empirical tests based on these feedbacks is immense (Kulmatiski et al., 2008; Van der Putten et al., 2013) but most empirical responses focus on plant biomass. Utilizing this method to focus on PSF *training and test* rounds on specific plant traits, such as flowering, root architecture, or chemistry, is much rarer. Even when focusing on biomass, these designs consistently show context dependency, including from competition, edaphic properties, and climate (Beals et al., 2020; Hassan et al., 2022; Cheng et al., 2025). Utilizing PSF systems for plant traits like flowering may quickly show that generational feedback to flowering traits also depends on environmental context. In understanding these GxE interactions that play out through filtering of plant microbiomes, we may discover common contexts where microbiomes or genotypic effect dominate plant traits. Microbiomes already appear to serve as an important pathway in the genetic effects of heterosis in ways that may be paralleled in specific aboveground traits (Wagner, 2021; Wagner et al., 2021). Future work on genotype and environment-specific feedbacks through the microbiome can fill this critical knowledge gap for key fitness traits like flowering.

## Acknowledgements

This work was made possible through a University of Kansas General Research Fund (GRF) grant to BAS and JKK from the University of Kansas. BAS and PMH were partially supported by NSF Award No. OIA-1656006 and NSF DBI-2120153. We thank Alyssa Larson for her preliminary work on these flowering tests. JKK acknowledges support from NSF award MCB-1940785. We thank Luis Madrigal Roca for writing the R script to implement elastic net regression.

## Author Contributions

BAS and JKK designed the experiment and obtained funding. PMH, JKK, and BAS carried out experiments and data analysis. BAS and JKK drafted and edited the manuscript.

## Data Availability Statement

Trait measurement data is available at ____. 16S sequencing data from Round 1 training are deposited in NCBI’s Short Read Archive under PRJA XXXXXXX.

## SUPPLEMENTAL INFORMATION

**Table S1:** ANOVA tables of treatment effects on flowering traits from REML in linear mixed effect models.

*Days to Flower:*
|  | Sum Sq | Mean Sq | NumDF | DenDF | F value | Pr(>F) |  |
| --- | --- | --- | --- | --- | --- | --- | --- |
| Rd2.Geno | 1988.79 | 1988.79 | 1 | 65.164 | 55.1926 | 2.97E-10 | *** |
| Soil | 290.89 | 290.89 | 1 | 67.622 | 8.0729 | 0.005933 | ** |
| MatGeno | 127.37 | 127.37 | 1 | 33.721 | 3.5349 | 0.068753 | . |
| Rd2.Geno:Soil | 49.21 | 49.21 | 1 | 65.658 | 1.3658 | 0.246769 |  |
| Rd2.Geno:MatGeno | 8.85 | 8.85 | 1 | 65.164 | 0.2457 | 0.62176 |  |
| Soil:MatGeno | 191.19 | 191.19 | 1 | 67.622 | 5.3059 | 0.024334 | * |
| Rd2.Geno:Soil:MatGeno | 4 | 4 | 1 | 65.658 | 0.111 | 0.74007 |  |

*Corolla Width:*
|  | Sum Sq | Mean Sq | NumDF | DenDF | F value | Pr(>F) |  |
| --- | --- | --- | --- | --- | --- | --- | --- |
| Rd2.Geno | 6568.3 | 6568.3 | 1 | 75 | 66.5205 | 6.02E-12 | *** |
| Soil | 621.2 | 621.2 | 1 | 75 | 6.291 | 0.0143 | * |
| MatGeno | 151.9 | 151.9 | 1 | 75 | 1.5388 | 0.2187 |  |
| Rd2.Geno:Soil | 1.8 | 1.8 | 1 | 75 | 0.0186 | 0.8919 |  |
| Rd2.Geno:MatGeno | 177.5 | 177.5 | 1 | 75 | 1.7973 | 0.1841 |  |
| Soil:MatGeno | 100.2 | 100.2 | 1 | 75 | 1.0146 | 0.317 |  |
| Rd2.Geno:Soil:MatGeno | 37.2 | 37.2 | 1 | 75 | 0.377 | 0.541 |  |

*Pistil Length:*
|  | Sum Sq | Mean Sq | NumDF | DenDF | F value | Pr(>F) |  |
| --- | --- | --- | --- | --- | --- | --- | --- |
| Rd2.Geno | 683.58 | 683.58 | 1 | 76 | 47.4522 | 1.42E-09 | *** |
| Soil | 18.74 | 18.74 | 1 | 76 | 1.3012 | 0.257579 |  |
| MatGeno | 27.25 | 27.25 | 1 | 76 | 1.8917 | 0.173052 |  |
| Rd2.Geno:Soil | 148.82 | 148.82 | 1 | 76 | 10.3309 | 0.001921 | ** |
| Rd2.Geno:MatGeno | 2.95 | 2.95 | 1 | 76 | 0.2049 | 0.65211 |  |
| Soil:MatGeno | 18.76 | 18.76 | 1 | 76 | 1.3021 | 0.257407 |  |
| Rd2.Geno:Soil:MatGeno | 65.24 | 65.24 | 1 | 76 | 4.5291 | 0.036565 | * |

*Biomass:*
|  | Sum Sq | Mean Sq | NumDF | DenDF | F value | Pr(>F) |  |
| --- | --- | --- | --- | --- | --- | --- | --- |
| Rd2.Geno | 46.365 | 46.365 | 1 | 75.48 | 105.7384 | 5.24E-16 | *** |
| Soil | 0.006 | 0.006 | 1 | 74.918 | 0.0147 | 0.90374 |  |
| MatGeno | 0.443 | 0.443 | 1 | 39.418 | 1.0105 | 0.32093 |  |
| Rd2.Geno:Soil | 0.078 | 0.078 | 1 | 74.204 | 0.1787 | 0.6737 |  |
| Rd2.Geno:MatGeno | 0.145 | 0.145 | 1 | 75.48 | 0.3308 | 0.56691 |  |
| Soil:MatGeno | 1.358 | 1.358 | 1 | 74.918 | 3.0966 | 0.08254 | . |
| Rd2.Geno:Soil:MatGeno | 0.677 | 0.677 | 1 | 74.204 | 1.5429 | 0.2181 |  |

**Table S2:** Elastic regression top five hits based on coefficients for each Round 2 trait. AM_I_IM62 represents the coefficient for the test plant genotype on the trait in that round.

| **<u>Taxonomy Match - Level</u>** | **<u>Coefficient</u>** | **<u>Response</u>** |
| --- | --- | --- |
| Aquicella - Genus | 1.32 | Biomass |
| Vicinamibacterales - Order | -0.475 | Biomass |
| AM-I-IM62 | 0.449 | Biomass |
| Solirubrobacter - Genus | 0.427 | Biomass |
| Acetobacteraceae - Family | 0.381 | Biomass |
| Litorilinea - Genus | 7.71 | FL.PISTIL |
| Candidatus Udaeobacter - Genus | 6.09 | FL.PISTIL |
| Planctomycetota OM190 - Class | 5.49 | FL.PISTIL |
| Candidatus Xiphinematobacter - Genus | -4.77 | FL.PISTIL |
| Phenylobacterium - Genus | -4.29 | FL.PISTIL |
| Blastocatellia 11-24 - Order | -25.3 | FL.W |
| Holophagaceae - Family | -19.9 | FL.W |
| Rokubacteriales - Order | -19.4 | FL.W |
| AM-I-IM62 | 13.6 | FL.W |
| Rubinisphaeraceae SH-PL14 - Genus | 12.2 | FL.W |
| Acetobacteraceae - Family | 16.6 | Time.to.Flow |
| AM-I-IM62 | 10.5 | Time.to.Flow |
| Tepidisphaerales WD2101 soil group - Family | 8.59 | Time.to.Flow |
| Solirubrobacteriales 67-14 - Family | 7.39 | Time.to.Flow |
| Caulobacteraceae - Family | 7.22 | Time.to.Flow |

**Table S3:**
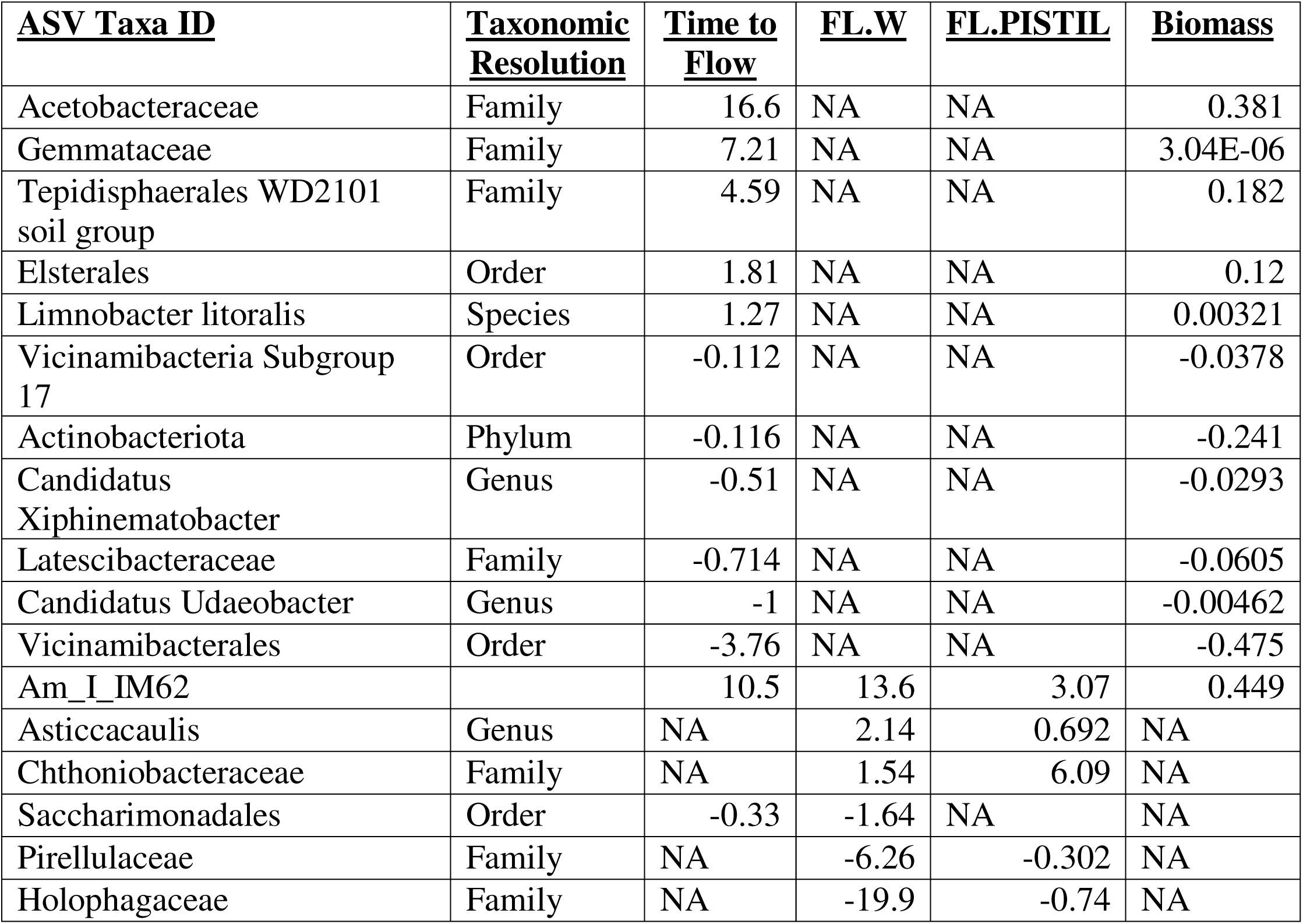
Elastic regression taxa of taxa with abundances significantly correlated with more than one trait in Round 2. AM_I_IM62 represents the coefficient for the test plant genotype on the trait in that round. Coefficients are only standardized within a trait, so magnitudes cannot be compared among traits.

**Supplemental Figure S1:**
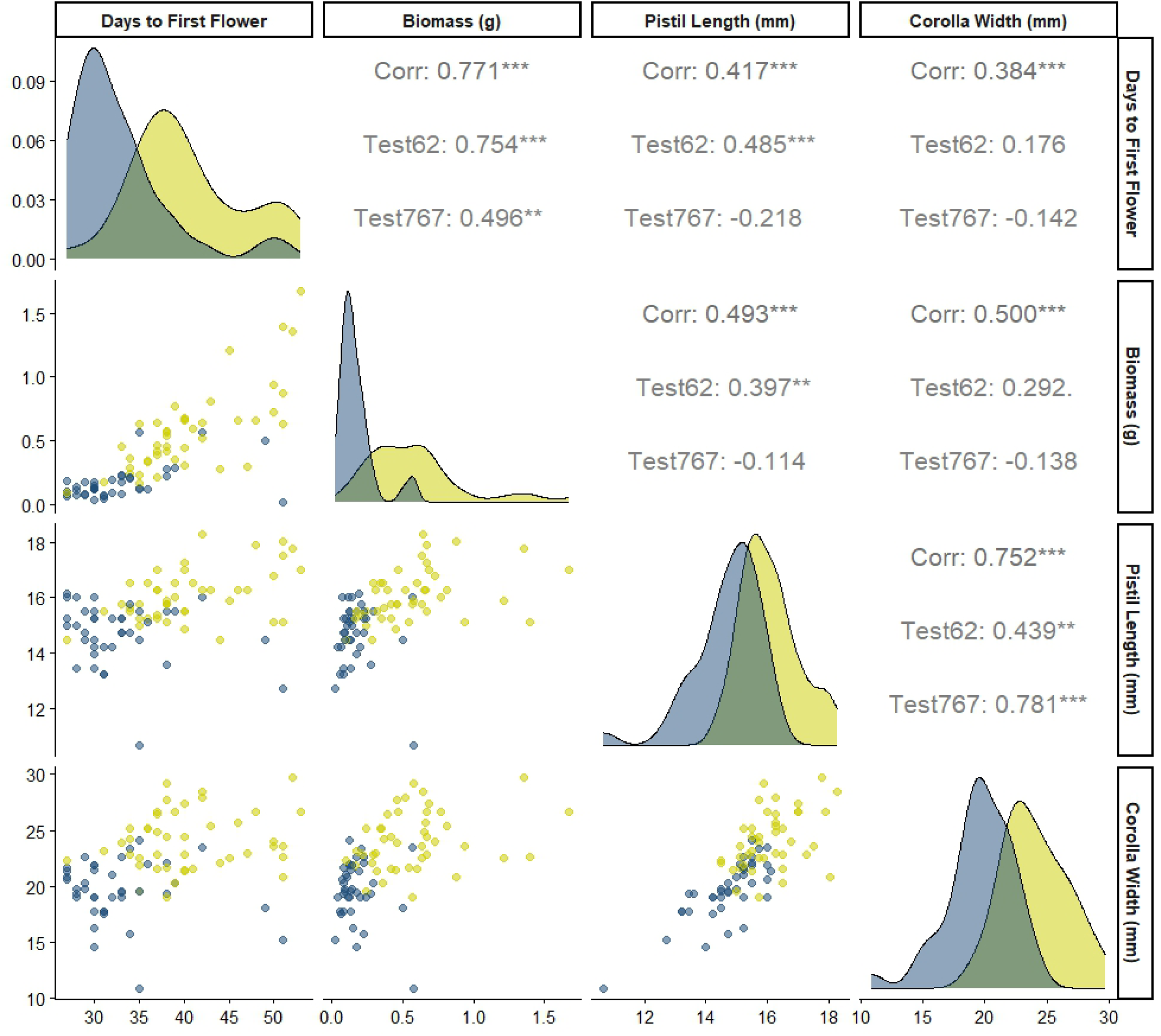
Pearson correlation tests among biomass and flowering traits overall and for a given test plant genotype.

**Supplemental Figure S2:**
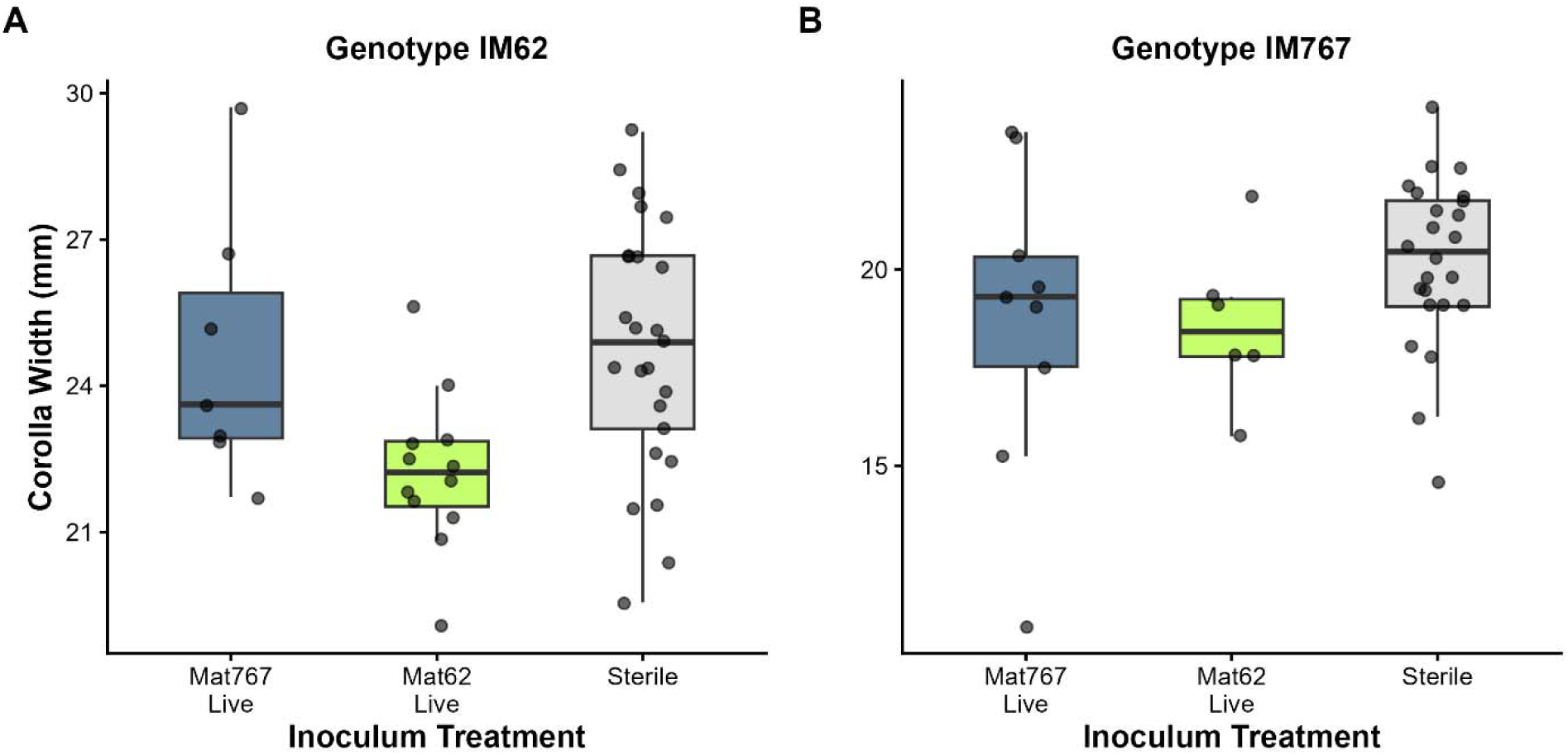
Corolla Width responses to test plant genotype, training round genotype of microbiome, or sterile inoculum. Inoculum is on the X-axis. Colors and notations are as in Figure 3.

**Supplemental Figure S3:**
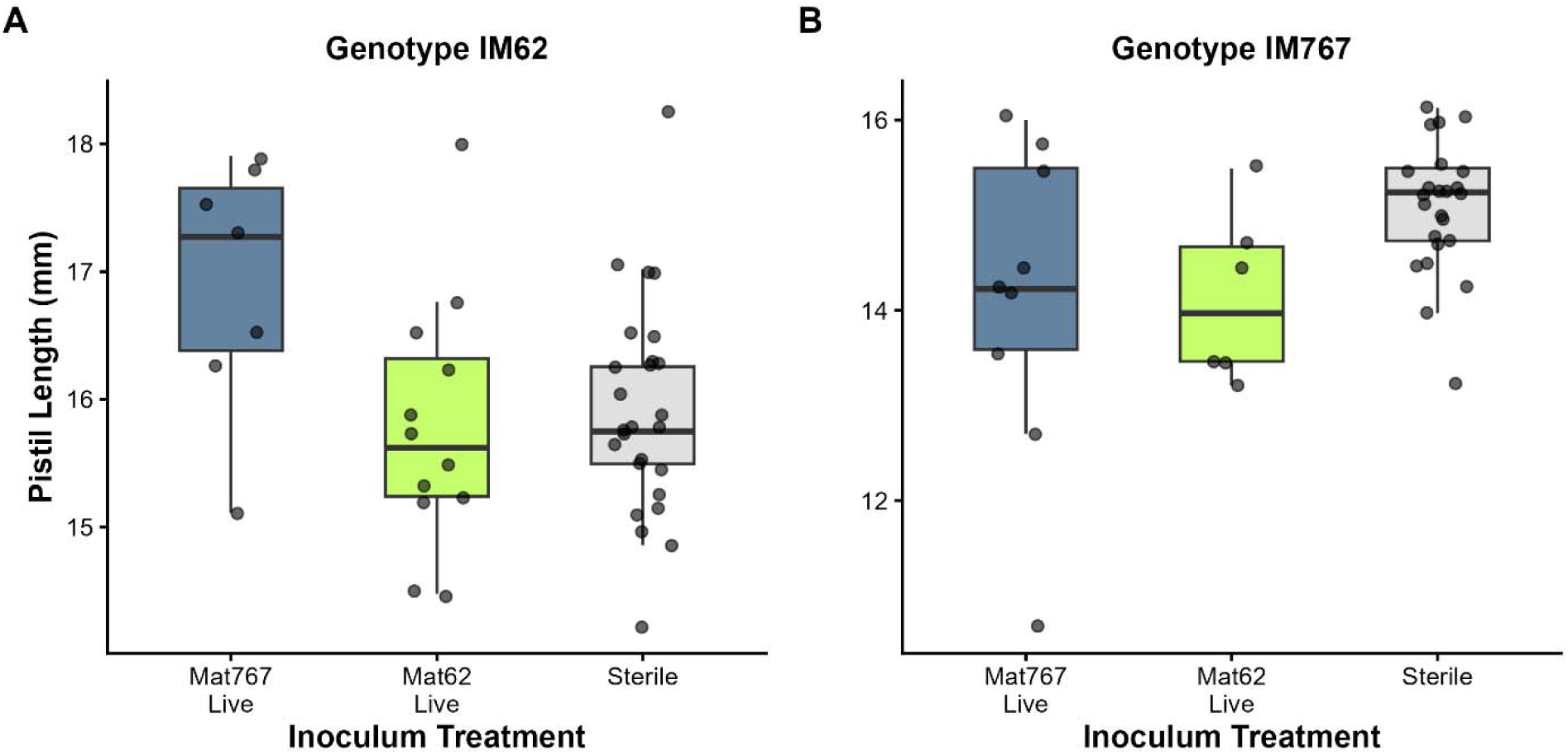
Pistil length responses to test plant genotype, training round genotype of microbiome or sterile inoculum. Inoculum is on the X-axis. Colors and notations are as in Figure 3.

**Supplemental Figure S4:**
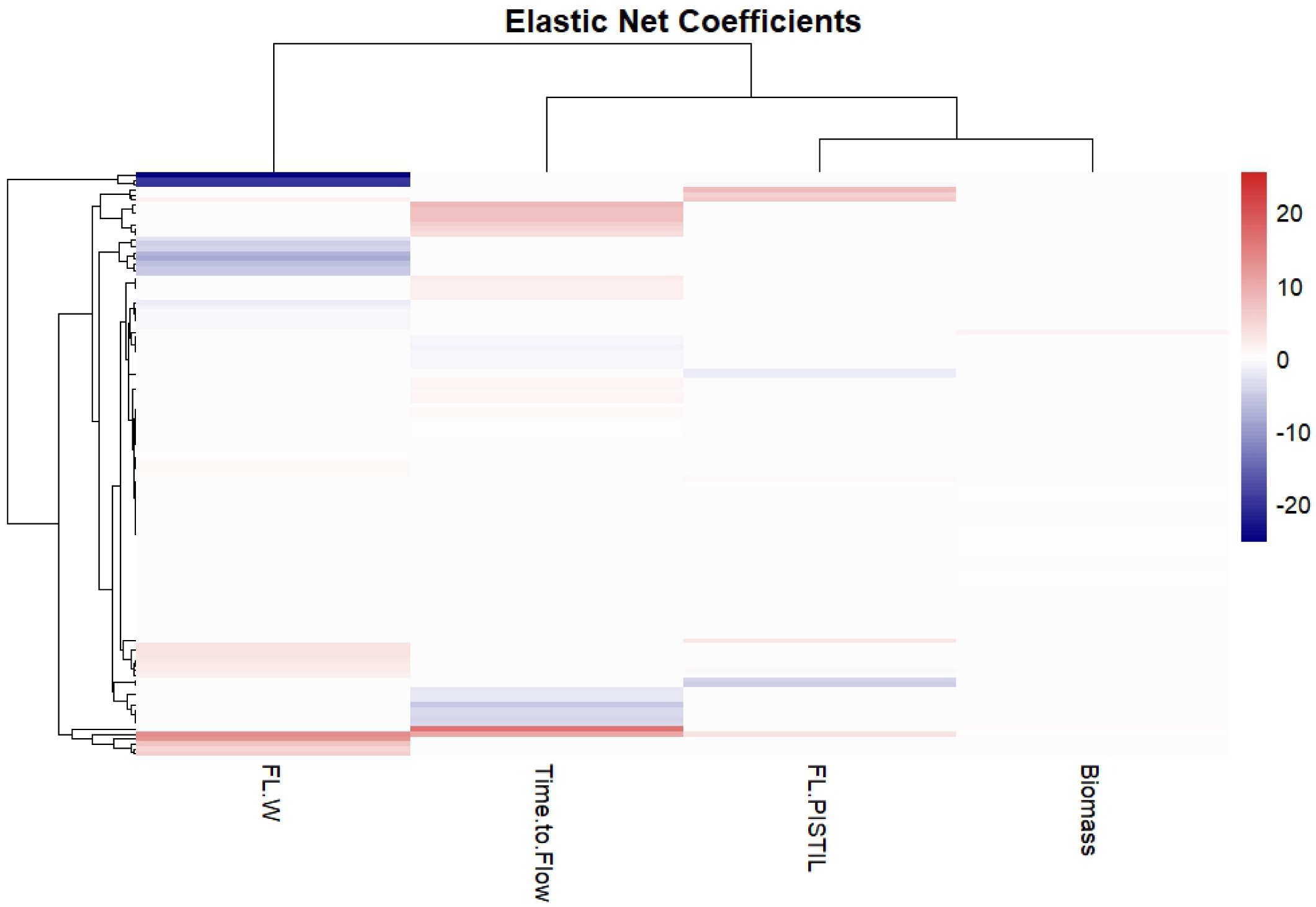
Heatmap of 16S ASV elements from training round associated with four traits in Round 2.

